# Phototroph-heterotroph interactions during growth and long-term starvation across *Prochlorococcus* and *Alteromonas* diversity

**DOI:** 10.1101/2021.10.26.465881

**Authors:** Osnat Weissberg, Dikla Aharonovich, Daniel Sher

**Author notes:** Dikla Aharonovich and Osnat Weissberg contributed equally to this study.

## Abstract

Microbial interactions such as those between phytoplankton and bacteria been studied intensively using specific model organisms, due to their potential impact on ecosystems and biogeochemistry. Yet, to what extent interactions differ between closely related organisms, or how these interactions change over time or culture conditions, remains unclear. Here, we characterize the interactions between five strains each of two globally abundant marine microorganisms, *Prochlorococcus* (a phototroph) and *Alteromonas* (a heterotroph), from the first encounter between individual strains and over more than a year of repeated cycles of exponential growth and long-term nitrogen starvation. *Prochlorococcus*-*Alteromonas* interactions had little effect on traditional growth parameters such as *Prochlorococcus* growth rate, maximal fluorescence or lag phase, affecting primarily the dynamics of culture decline, which we interpret as representing cell mortality and lysis. The shape of the *Prochlorococcus* decline curve and the carrying capacity of the co-cultures were determined by the phototroph and not the heterotroph strains involved. Comparing various mathematical models of culture mortality suggests that *Prochlorococcus* death rate increases over time in mono-cultures but decreases in co-cultures, with cells potentially becoming more resistant to stress. Our results demonstrate intra-species differences in ecologically-relevant co-culture outcomes. These include the recycling efficiency of N and whether the interactions are mutually synergistic or competitive. They also highlight the information-rich growth and death curves as a useful readout of the interaction phenotype.

**Significance Statement:** Interactions between phytoplankton and marine bacteria impact global ecosystems and biogeochemistry. Here, we explore how intra-species variability affects the interactions between *Prochlorococcus*, a globally abundant photosynthetic cyanobacetrium and *Alteromonas*, a heterotrophic bacterium that lives off and recycles organic matter. Under nitrogen starvation, *Prochlorococcus* growing alone increasingly accumulate damage and die, whereas in co-culture with *Alteromonas* they become increasingly resilient. The specific *Prochlorococcus* strain had a much larger effect on co-culture behavior than the *Alteromonas* strain, determining whether the interactions are mutually synergistic or potentially competitive. These results show how ecologically relevant outcomes of interactions may vary between closely related microorganisms, and highlight growth and death curves from laboratory (co)-cultures as information-rich views of microbial growth and death.

## Introduction

Interactions among microorganisms occur in every known ecosystem (recently reviewed by (1,2)). Detailed studies of the interactions between selected model organisms (often in laboratory co-cultures) have begun to reveal the diversity of molecular mechanisms whereby organisms interact with each other (2–4). However, it is currently unknown to what extent the studied interactions differ between organism pairs, growth stages or environmental conditions. For example, while broad-scale phylogenetic patterns are often observed in microbial interactions, closely related bacteria may differ in the way they interact with other organisms, likely as a result of the significant genetic diversity observed in many microbial clades (e.g. (5,6)). Additionally, the same pair of interacting organisms can synergize or compete depending on the composition of the culture media and the growth stage of (co)-culture (e.g. (7–9)). Finally, both the coarse-grained ecological classification of microbial interactions (e.g. positive/negative) and the high-resolution mechanistic view obtained using advanced physiology and ‘omics approaches are difficult to translate into quantitative, predictive models of organismal growth and decline (1,10,11).

Here, we explore to what extent intra-clade diversity affects the outcome of microbial interactions, using growth curves as an information-rich view of microbial growth and mortality. Growth curves can be divided into discrete phases (lag, exponential, stationary, decline and long-term stationary phases), and can be used to extract quantitative parameters such as growth rates, lag times, etc. (12,13). An extra layer of more subtle information may exist in the shapes of the growth curves, providing hints of important shifts in the physiology of the growing organisms, as classically demonstrated by Jacques Monod for diauxic growth in *E. coli* (14). Notably, while many studies of bacterial interactions focus on the exponential growth stage or on culture yield at a specific time-point (e.g. (15–17)), fewer studies look at the shape and dynamics of the decline phases, which can provide important hints regarding the effect of interactions on the process of microbial mortality (e.g. (18–20)).

Our model organisms are two globally abundant clades of marine bacteria: a cyanobacterial primary producer (*Prochlorococcus*) and a heterotrophic γ-proteobacterium (*Alteromonas*). Interactions between marine phototrophs (phytoplankton, including cyanobacteria) and heterotrophic bacteria have been studied intensively, as phytoplankton are responsible for about one-half of the photosynthesis on Earth (e.g. (21–25)). Thus, phytoplankton-bacteria interactions may strongly affect community structure and function on scales from microns to thousands of kilometers (26,27). Our model primary producer, *Prochlorococcus*, is found throughout the euphotic zone, the sunlit upper portion, of the oligotrophic (nutrient-poor) ocean. There are multiple *Prochlorococcus* clades, broadly partitioned into high-light (HL) and low light (LL) adapted ecotypes, which differ in their photosynthetic parameters and occupy different niches in the ocean (e.g. surface vs deep water, reviewed by (28)). Strains differ also in traits such as the capacity to utilize different forms of inorganic nutrients and organic matter, as well as in their interactions with heterotrophic bacteria and phage. *Alteromonas* is a clade of free-living marine bacteria, which are also partitioned into surface and deep groups (*A. macleodii* and *A. mediterranea*, respectively) (29). *Alteromonas* strains also exhibit diverse capabilities to utilize carbohydrates, to acquire iron, and in motility (30). Interactions between individual strains of *Prochlorococcus* and *Alteromonas* have been characterized in some detail (12,27,31–35). While the phenotype and gene expression patterns during interactions vary between strains, this variability has not been explored systematically (12,32,36). Notably, strain- and condition-dependent phytoplankton-heterotroph interactions are observed also in other systems, including *Synechococcus*, a close relative of *Prochlorococcus* (18,37,38), as well as eukaryotic microalgae (e.g. coccolithophores and diatoms, (7–9,39)).

We characterized the interactions between five strains each of *Prochlorococcus* and *Alteromonas*, from the first encounter between previously-axenic strains (i.e., grown in mono-culture) and across ∼1.2 years of co-culture (25 phototroph-heterotroph combinations, Figure 1A, Supplementary Figure S1). The culturing period spanned multiple cycles of exponential growth, culture decline and long-term nitrogen starvation (33). Nitrogen limitation occurs across wide swaths of the global ocean, and affects a significant fraction of the *Prochlorococcus* diversity (40,41). We focused our analysis on *Prochlorococcus* growth and decline, which is easily measured using bulk culture fluorescence in a non-invasive manner (e.g. (12,42)). Using this dataset of 429 growth curves, as well as associated cell counts, we ask: i) How do the interactions between *Prochlorococcus* and *Alteromonas* vary across the diversity of each organisms? ii) Do the interactions change over time (i.e. do the organisms adapt to “living together”)? iii) When, during the life-cycle of a *Prochlorococcus* batch culture, do microbial interactions have the largest impact on growth, death, and overall culture carrying capacity, and can this impact be quantified?

**Figure 1:**
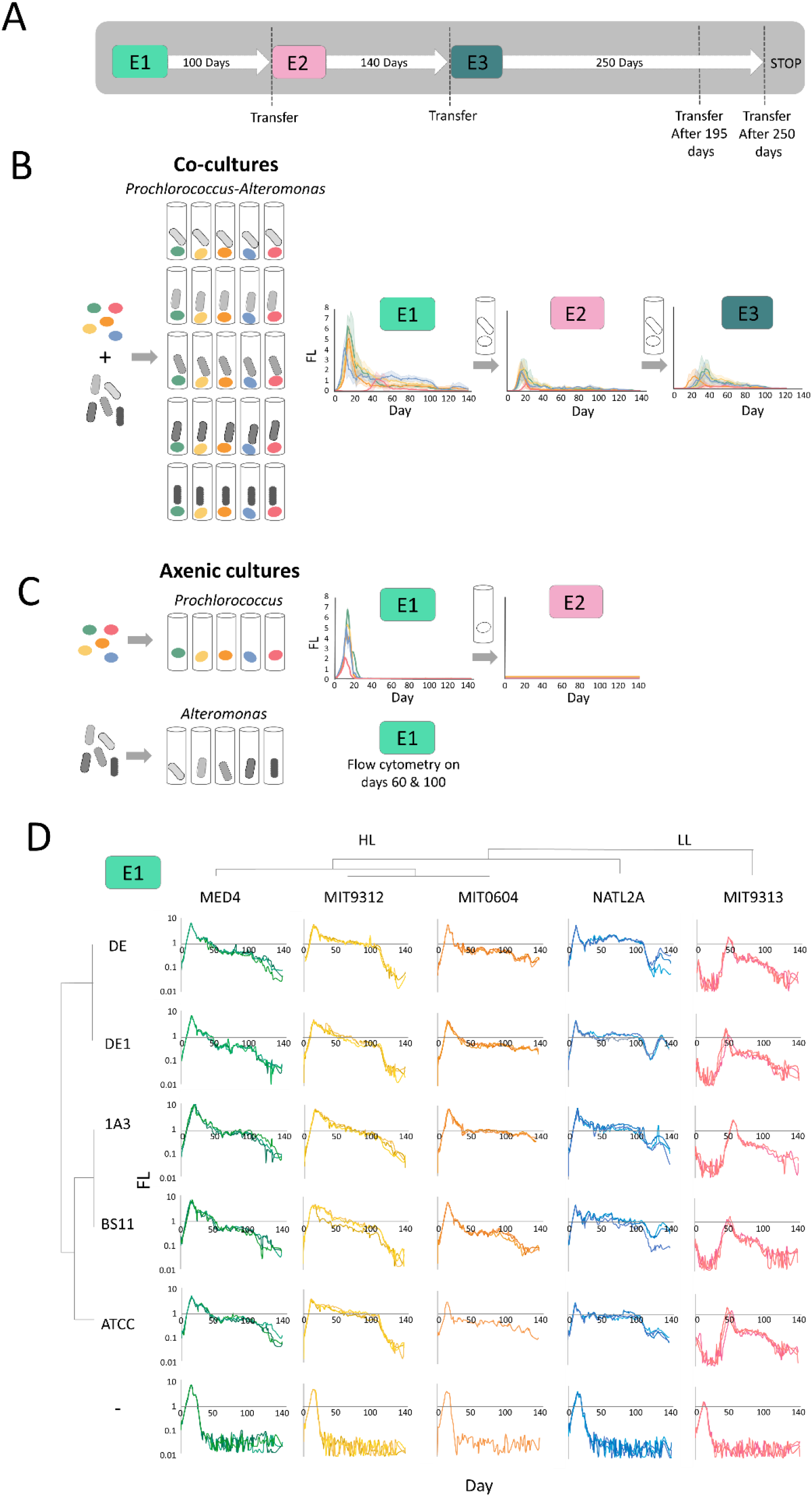
Experimental designs and overview of the dynamics of *Prochlorococcus*-*Alteromonas* co-cultures from first encounter and over multiple transfers. A. Schematic illustration of the experimental design. 1 ml from Experiment E1 was transferred into 20 ml fresh media after 100 days, starting experiment E2. Experiment E2 was similarly transferred into fresh media after 140 days, starting experiment E3. Additional experiments replicating these transfers are described in Supplementary figure S1. B. Overview of the growth curves of the 25 *Prochlorococcus-Alteromonas* co-cultures over three transfers spanning ∼1.2 years (E1, E2 and E3). Results show mean + standard error from biological triplicates, colored by *Prochlorococcus* strain as in panel D. C. Axenic *Prochlorococcus* grew exponentially in E1 but failed to grow when transferred into fresh media after 60 days. Axenic *Alteromonas* cultures were counted after 60 and 100 days, as their growth cannot be monitored sensitively and non-invasively using fluorescence (optical density is low at these cell numbers). D. High reproducibility and strain-specific dynamics of the initial contact between *Prochlorococcus* and *Alteromonas* strains (E1). Three biological replicates for each mono-culture and co-culture are shown. Note that the Y axis is linear in panels B, C and logarithmic in panel D.

## Results

### All *Alteromonas* strains support long-term survival of *Prochlorococcus* under N starvation

*Prochlorococcus*, and to some extent *Synechococcus*, have previously been shown depend on co-occurring heterotrophic bacteria to survive various types of stress, including nitrogen starvation (33,34,42,43). At the first encounter between previously-axenic *Prochlorococcus* and *Alteromonas* (E1), all co-cultures and axenic controls grew exponentially (Figure 1B, C). However, all axenic cultures showed a rapid and mostly monotonic decrease in fluorescence starting shortly after the cultures stopped growing, reaching levels below the limit of detection after ∼20-30 days. None of the axenic *Prochlorococcus* cultures were able to re-grow when transferred into fresh media after 60 days (Figure 1C). In contrast, the decline of co-cultures rapidly slowed, and in some cases was interrupted by an extended “plateau” or second growth stage (Figure 1B). Across multiple experiments, 92% of the co-cultures contained living *Prochlorococcus* cells for at least 140 days, meaning that they could be revived by transfer into fresh media. Thus, the ability of *Alteromonas* to support long-term N starvation in *Prochlorococcus* was conserved in all analyzed strains.

It has previously been shown that *Prochlorococcus* MIT9313 is initially inhibited by co-culture with *Alteromonas* HOT1A3, while *Prochlorococcus* MED4 is not (12,32). This “delayed growth” phenotype was observed here too, was specific to MIT9313 (not observed in other *Prochlorococcus* strains) and occurred with all *Alteromonas* strains tested (Figure 1D). MIT9313 belongs to the low-light adapted clade IV (LLIV), which are relatively distant from other *Prochlorococcus* strains and differ from them in multiple physiological aspects including the structure of their cell wall (44), the use of different (and nitrogen-containing) compatible solutes (45) and the production of multiple peptide secondary metabolites (lanthipeptides,(46,47)). LLIV cells also have larger genomes, and are predicted to take up a higher diversity of organic compounds such as sugars and amino acids (48). It is intriguing that this strain, which has higher predicted metabolic and regulatory flexibilities (49), is the only one initially inhibited in co-culture with *Alteromonas*.

### Differences in co-culture phenotype are related to *Prochlorococcus* and not *Alteromonas* strains and occur primarily during the decline stage

While co-culture with all *Alteromonas* strains had a major effect on *Prochlorococcus* viability during long-term starvation, there was no significant effect of co-culture on traditional metrics of growth such as maximal growth rate, maximal fluorescence and lag phase (with the exception of the previously-described inhibition of MIT9313, Figure 2A,B,C). However, a visual inspection of the growth curves in Figure 1D suggested subtle yet consistent differences in the shape of the growth curve, and especially the decline phase, between the different *Prochlorococcus* strains in the co-cultures. To test this, we used the growth curves as input for a Principal Component Analysis (PCA), revealing that the growth curves from each *Prochlorococcus* strain clustered together, regardless of which *Alteromonas* strain they were co-cultured with (Figure 2D). The growth curves of all high-light adapted strains (MED4, MIT9312 and MIT0604) were relatively similar, the low-light I strain NATL2A was somewhat distinct, and the low-light IV strain MIT9313 was a clear outlier (Figure 1D), consistent with this strain being the only one initially inhibited in all co-cultures. Random Forest Classification (a supervised machine learning algorithm) supported the observation that the growth curve shapes were affected more by the *Prochlorococcus* rather than *Alteromonas* strains, and also confirmed the visual observation that most of the features differentiating between *Prochlorococcus* strains occurred during culture decline (Supplementary Figure S2, Supplementary Text S1). Thus, co-culture with *Alteromonas* affects the decline stage of *Prochlorococcus* in co-culture in a way that differs between *Prochlorococcus* but not *Alteromonas* strains.

**Figure 2:**
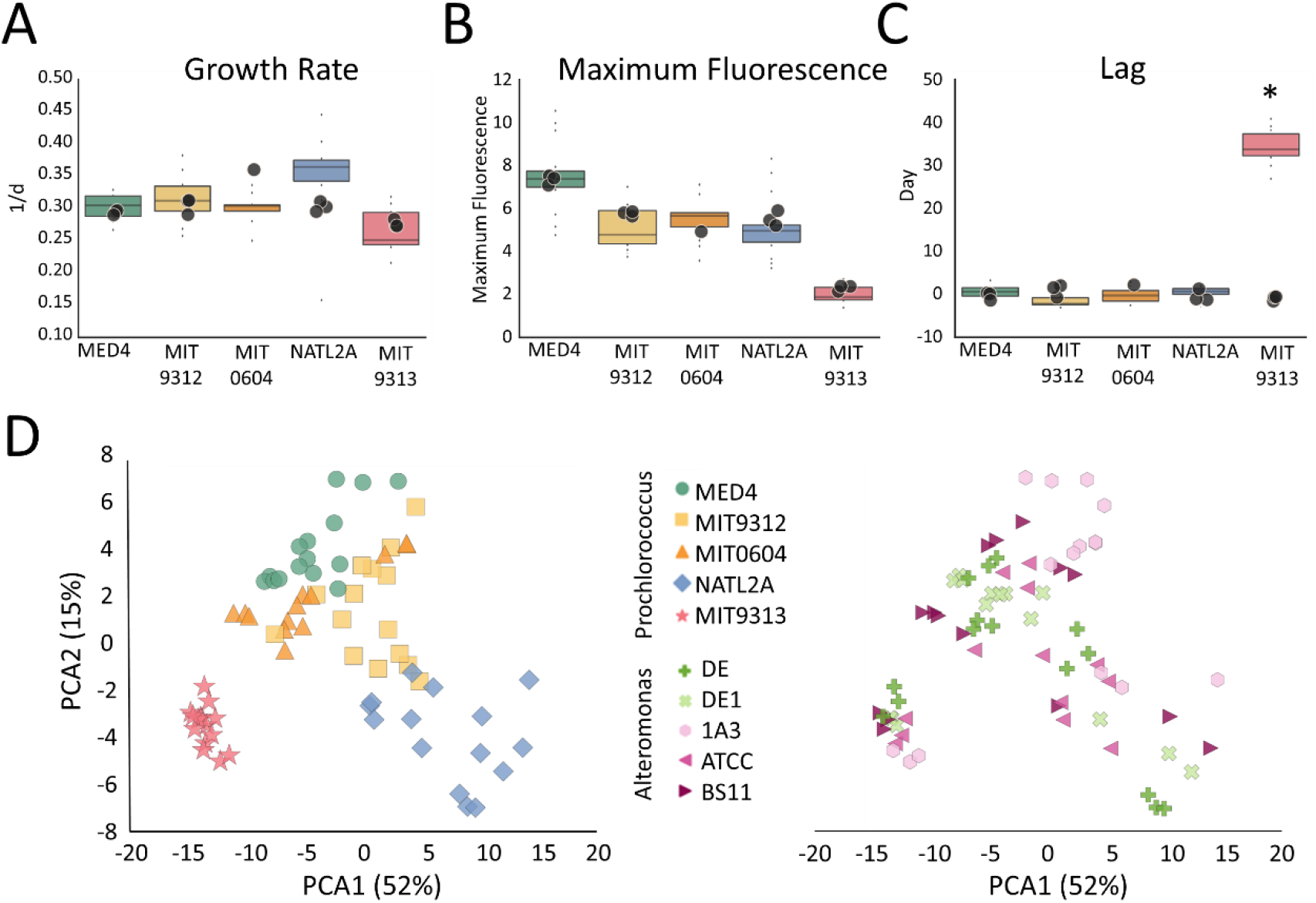
Growth analysis and principal component analysis (PCA) of the growth curves from all co-cultures during 140 days of E1. A. Growth rate, Maximum fluorescence and Lag phase during E1. Box-plots represent mean and 75^th^ percentile of co-cultures, circles represent measurements of individual cultures of the axenic controls. The only significant difference between axenic and co-cultures is in the length of the lag phase for MIT9313 (Bonferroni corrected ANOVA, p < 0.001. B. PCA ordination colored by *Prochlorococcus* (left) and by *Alteromonas* (right) strains. The growth curves cluster by *Prochlorococcus* strain (Adonis test, F(4,68) = 42.3, p = 0.001) and only marginally by *Alteromonas* strain (Adonis test, F(4,68) = 2.29, p = 0.017)

We next asked whether the phenotypes of interaction, which were observed when high cell densities of *Prochlorococcus* and *Alteromonas* interacted for the first time (E1), were maintained after the cells had grown together in co culture for extended periods. We therefore continued to transfer the co-cultures into fresh media over multiple additional transfers, performed 40-200 days after the initial inoculations. In total, FL measurements are available for a cumulative period of 380 days which the cells spent in co-culture (Figure 1A, Supplementary Figure S1). The ability of *Prochlorococcus* to survive long-term N starvation, the clustering of the growth curves by *Prochlorococcus* but not *Alteromonas* strains, and the results of the Random Forest Classification, were all reproduced in subsequent transfers (Figure 1B, Supplementary figures S2, S3, S4, Supplementary Text S2). These observations are thus robust to the cumulative time the organisms have been interacting and the cell densities of both organisms when transferred to new media (see below).

### Differences in the carrying capacity suggest different modes of interaction

While *Alteromonas* clearly support *Prochlorococcus*, by enabling it to survive long-term N starvation, is the reciprocal interaction also synergistic? Do *Prochlorococcus* enhance the growth of *Alteromonas*, and does the interaction affect the overall carrying capacity of the system, defined here as the ability to efficiently utilize the limiting resource (nitrogen)? To answer these questions, we used the flow cytometry cell counts of *Prochlorococcus* and *Alteromonas* on days 60, 100 and 140 to infer the nitrogen (N) biomass of each population grown alone or in co-culture (Figure 3A, Supplementary Table S1, see supplementary text S3 for the calculations and caveats).

**Figure 3:**
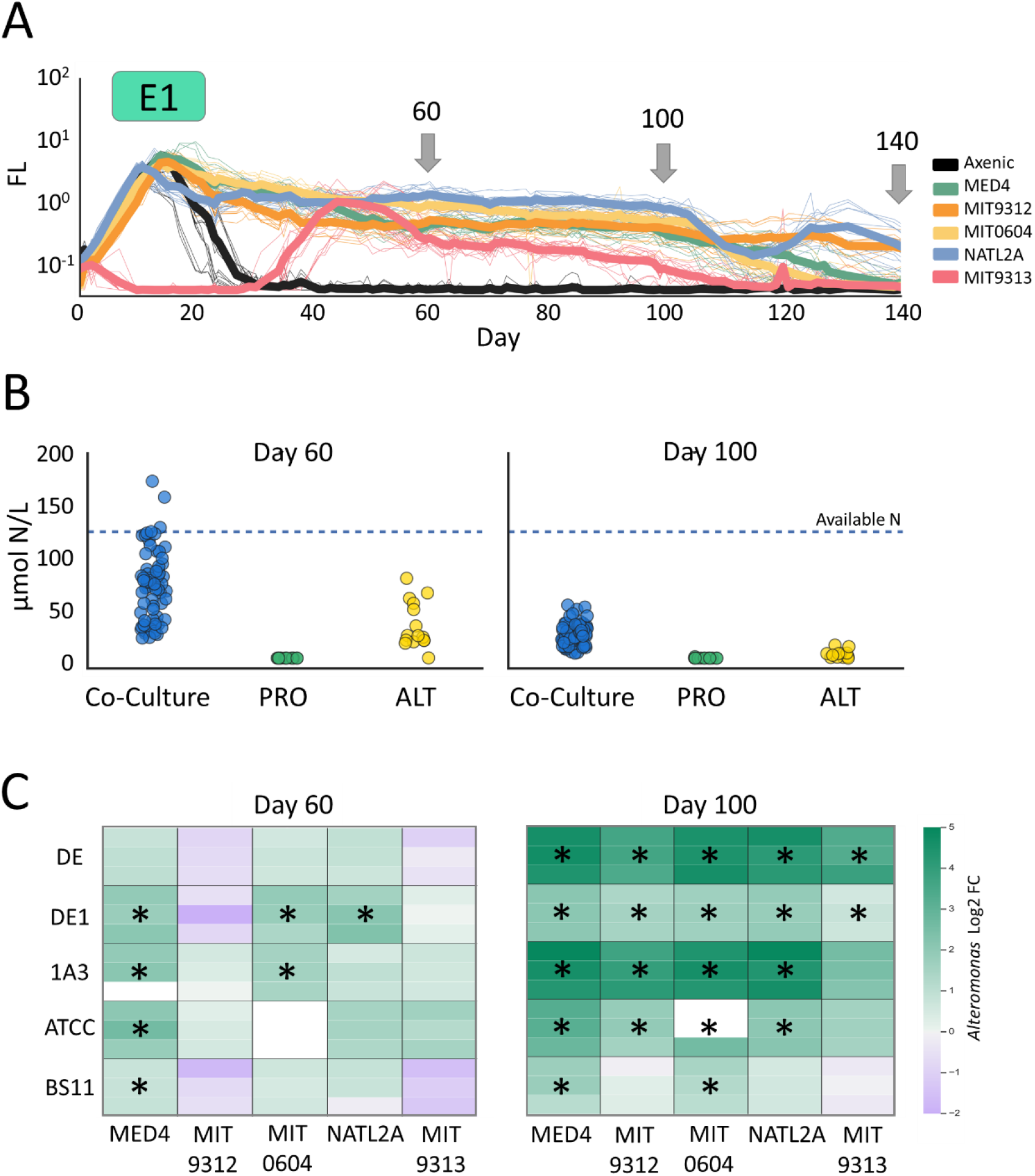
Carrying capacity and type of interactions. A. Growth curves of experiment E1, arrows showing the days where cell numbers were counted by flow cytometry (see also Supplementary Table S1). Axenic curves shown are from all *Prochlorococcus* strains. B. Total calculated N biomass of the cultures in E1 on days 60 and 100 in μmol N/L. Dashed line indicates total available N in co-cultures. C. Log2 Fold Change (FC) of *Alteromonas* N biomass relative to axenic *Alteromonas* controls on days 60 and 100. Asterisks indicate statistically significant FC (Bonferroni corrected ANOVA p < 0.05). ALT: *Alteromonas* PRO: *Prochlorococcus*

The overall carrying capacity of the system was higher than the axenic *Alteromonas* cultures, and much higher than the axenic *Prochlorococcus* (Figure 3B). On day 60, the mean carrying capacity of the co-cultures was 2-3 times higher than that of the Axenic *Alteromonas* (69±35 compared with 32±22 μmol N/L), suggesting that the heterotroph benefited from carbon fixed by the phototrophic *Prochlorococcus* partner. Indeed, most of this cellular N was found in the *Alteromonas* cells (76±13%). The ability of axenic *Alteromonas* to survive in the absence of organic matter from *Prochlorococcus* is not surprising, as an *Alteromonas* strain distantly related to the ones studied here, AltSIO, can utilize a large fraction of the labile organic material found in natural seawater used to make the growth media (50). In contrast, in axenic *Prochlorococcus* cultures only a small fraction of N in the system was found in cell biomass (∼0.01 μmol N/L). This likely reflects the inability of all *Prochlorococcus* strains to recycle organic nitrogen lost due to exudation or cell lysis (Figure 3B).

However, mutual synergism was not observed across all strain combinations. While some *Prochlorococcus* strains (MED4, MIT0604, and NATL2A) supported significantly higher *Alteromonas* N biomass compared to the axenic control (log2FC 1.3±0.6), co-cultures with MIT9312 and MIT9313 resulted in similar or lower *Alteromonas* biomass (log2FC -0.2±0.9) (Figure 3C). Therefore, on day 60, some of the interactions were mutually synergistic whereas in other cases *Prochlorococcus* do not support *Alteromonas* and may even compete with it. Notably, the two “non-mutually-synergistic” *Prochlorococcus* strains belong to different ecotypes but were isolated from the same drop of water from the Gulf Stream (51). In all co-cultures the *Prochlorococcus* population benefited from the presence of *Alteromonas* (log2FC 10±4).

In contrast to day 60, after 100 days essentially all of the interactions were mutually synergistic, with *Alteromonas* supporting the growth of all *Prochlorococcus* strains (log2FC 10±1) and *Prochlorococcus* increasing the *Alteromonas* biomass in all strains with the exception of BS11 (log2FC 3±1.5) (figure 3C). This suggests that the mode of interaction (synergist vs competition) may change temporally during the extended period of N starvation.

On day 140 the carrying capacity of the co-cultures declined further (only 1% of the N was in biomass). This suggests that the system is not in steady state, with a slow yet constant reduction in carrying capacity. We speculate that this is driven by the loss of bioavailable N from the system (i.e., most of the nitrogen is in a recalcitrant form that cannot be utilized by either partner). This is further supported by the observation that while the co-cultures after 140 days were still alive and could be transferred to new media. In a subsequent experiment, only 16/30 cultures could be transferred after 195 days, and only 3/75 cultures could be transferred after 250 days (Supplementary Figure S5). Notably, MIT0604 (HLII) was the strain most likely to survive transfer after these extended periods and was also the most abundant *Prochlorococcus* strain after 140 days (1.46±1 μmol N/L MIT0604 biomass vs 0.26±0.56 μmol N/L for all other strains). While we do not currently have an explanation for the higher survival of this strain, it is noteworthy that it is the only strain to utilize nitrate (52).

### Modeling the effect of co-culture on *Prochlorococcus* mortality

Given that the most striking effect of co-culture was on the decline phase of the co-cultures, we asked whether we could quantify and model the effect of *Alteromonas* on *Prochlorococcus* mortality. Importantly, while the growth of bacteria has been extensively studied and modelled, the decline of bacterial cultures is much less studied, and mortality is rarely represented in ecological or biogeochemical models of microbial dynamics (53). Bacterial mortality has, however, often been modelled in the context of food safety and genome evolution, using either mechanistic or descriptive approaches (53–57). We chose to focus on four of these previously-described models which are relatively simple and have a clear biological interpretation (Table 1). The Exponential model is the simplest and most commonly used one, where a constant portion of the population dies over time (58). The Bi-exponential model is slightly more complex, representing two separate subpopulations in the community, each with its own death rate (55). The Weibull model is probabilistic, modeling a heterogeneous population with a diverse stress tolerance (53,59), finally, the Harmonic model employs a quadratic rate of decline which is often associated with predator-prey interactions or cellular encounter rates (58). When fitting each of these models to the decline phase of the growth curves, the Weibull model stands out as it has a low error for both axenic and co-cultures (Table 1) as well as in consequent transfers (Supplementary Table 1, the biexponential model is a better fit for the co-cultures but does not represent well the axenic ones). Based on the Weibull model, and assuming that culture fluorescence is related to the number of non-lysed cells in the media, axenic *Prochlorococcus* cells die more than ten-fold faster than cells in co-culture (2-decimal reduction time, td_2_, is 12.58 ± 3.85 days for axenic cultures and 316 ± 337 days for co-cultures).

**Table 1:** Mathematical description and biological interpretation of four models used to describe bacterial mortality.

| Mathematical model<sup>1</sup> | Derivative | # Params<sup>2</sup> | Physiological Interpretation | Axenic RMSE <sup>3</sup> (n=13) | Co-culture RMSE (n=343) | Reference |
| --- | --- | --- | --- | --- | --- | --- |
| Exponential<br>$X = X_0 e^{-a t}$ | $\frac{dX}{dt} = -aX$ | 1 | Constant ratio of the population dies in each time point | $0.26 \pm 0.11$ | $0.29 \pm 0.21$ | (58) |
| Bi-Exponential<br>$X = X_0(f e^{-a_1 t} + (1 - f) e^{-a_2 t})$ | $\frac{dX}{dt} = -f a_1 X_1 - (1 - f) a_2 X_2$ | 3 | Two subpopulations with different persistence under stress and death rates | $0.26 \pm 0.11$ | $0.16 \pm 0.12$ | (55) |
| Harmonic<br>$X = X_0 \frac{1}{1+a t}$ | $\frac{dX}{dt} = -a X^2$ | 1 | Quadratic mortality rate – dependence on cell-cell encounters | $0.50 \pm 0.19$ | $0.19 \pm 0.13$ | (58) |
| Weibull<br>$X = X_0 e^{-\frac{t^n}{a}}$ | $\frac{dX}{dt} = -c t^{n-1} X^*$ | 2 | Probabilistic model with a heterogeneous distribution of stress tolerance | $0.14 \pm 0.08$ | $0.18 \pm 0.11$ | (59) |

<sup>1</sup> In the mathematical models, $t$ stands for time since the beginning of culture decline, $X$ and $X_0$ are the current and initial cell numbers respectively, and the rest of the variables are model parameters ( $a, f, a_1, a_2, n$ )
<sup>2</sup> # Params – number of free parameters
<sup>3</sup> RMSE - Root Mean Square Error. Supplementary Table S2 shows also the Bayesian Information Criterion (BIC), which is less intuitive but takes into account also the number of model parameters.

In the Weibull model, the ‘shape parameter’ (n) represents the change over time in the susceptibility of the bacterial community to stress. A shape parameter above one represents an increasing probability that cells will die as time increases (e.g. due to the accumulation of damage), whereas a shape parameter below one suggests that, as the culture declines, the cells become more resistant to damage. Axenic cultures have high mean shape value of 2.1 ± 0.9, suggesting an accumulation of cell damage leading to increasing death rate (Figure 4A). In contrast, the mean shape value of co-cultures is significantly lower and below 1 (0.4 ± 0.2, student t-test, p< 0.001), suggesting that during N starvation in co-culture the *Prochlorococcus* cells are acclimating over time to the nutrient stress conditions.

**Figure 4:**
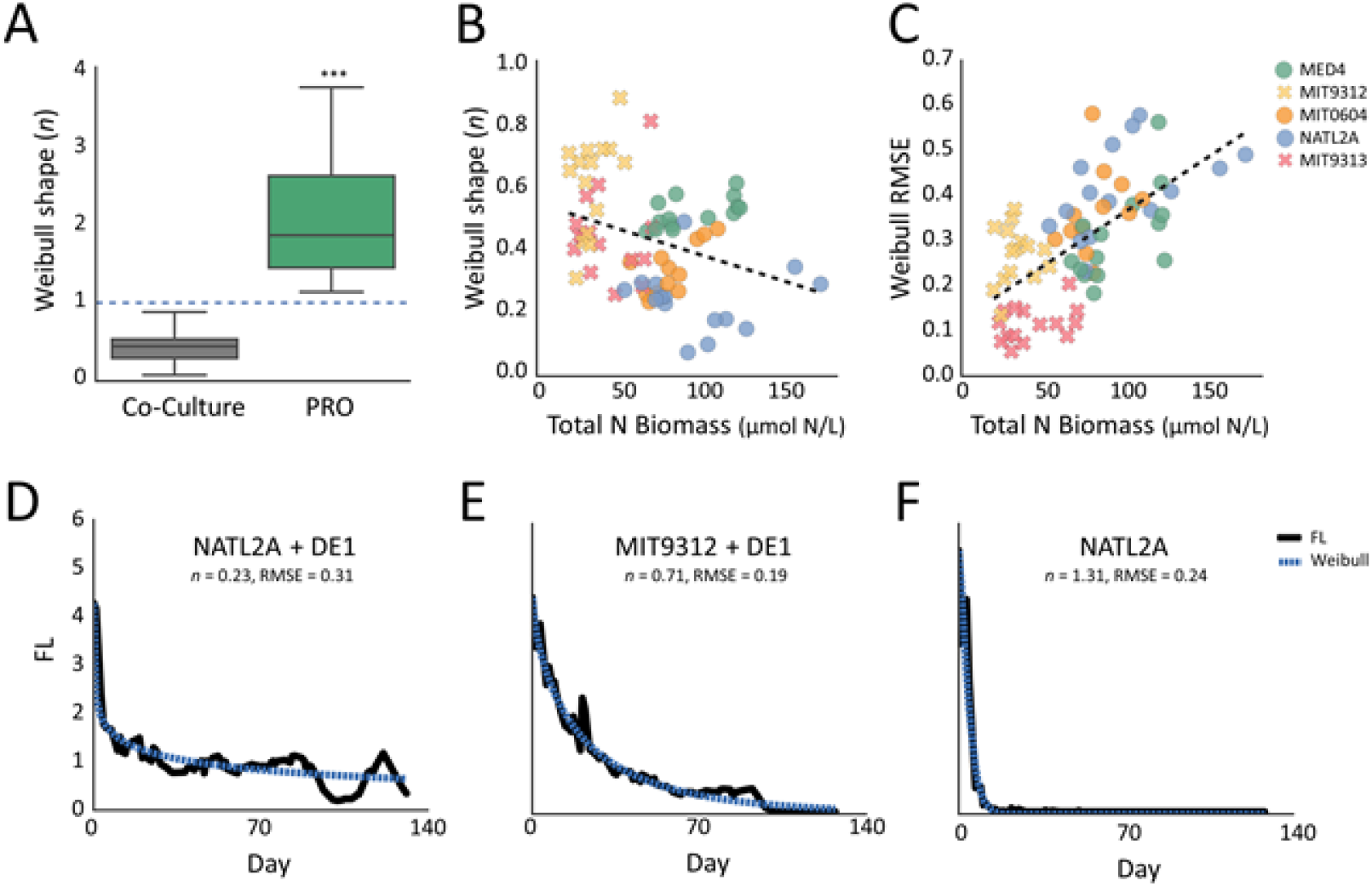
Weibull modeling of long-term starvation. A. Weibull shape (*n*) in Axenic *Prochlorococcus* (PRO) and in Co-cultures. In all axenic samples *n* > 1, in most co-cultures *n* < 1 (students t-test, p< 0.001). B. Scatter plot showing the reverse correlation between the total N biomass of the co-cultures on day 60 and Weibull shape (*n*). Pearson r = -0.33, p=5e-3. Circles represent co-cultures with mutual synergistic interactions (i.e. *Prochlorococcus* strains MED4, MIT0604 and NATL2A), X represent potential competitive interactions (strains MIT9312, MIT9313). C. Scatter plot showing the correlation between the total N biomass of the co-cultures on day 60 and Weibull root mean square error (RMSE). Pearson r = 0.64, p < 0.001. For the correlations with total N biomass on day 100 see Supplementary Figure S6. D-F. Weibull model fit for selected decline curves. D. Mutually synergistic co-culture of *Prochlorococcus* NATL2A and *Alteromonas* DE1. E. Potentially competitive co-culture of *Prochlorococcus* MIT9312 and *Alteromonas* DE1. F. Axenic *Prochlorococcus* NATL2A

While the molecular and physiological mechanisms of *Prochlorococcus* adaptation are currently unclear, the Weibull shape parameter decreases as the total N in cellular biomass increases, suggesting that the *Prochlorococcus* acclimation process is related to the ability to recycle N between the specific *Prochlorococcus* and *Alteromonas* strains in co-culture (Figure 4B). Thus, the rate of acclimation is higher in the co-cultures supporting high N biomass and mutually synergistic interaction (NATL2A, MED4 and MIT0604) compared to MIT9312 and MIT9313, where *Alteromonas* do not gain from the interaction (0.36±0.14 vs 0.52±0.16, t-test p < 0.001).

While the Weibull model is useful in quantifying mortality rates and raising the hypothesis that *Prochlorococcus* cells are acclimating to starvation over time, none of the tested models was able to fully capture the intricate dynamics of culture decline (Figure 4C, D). In most mutually synergistic co-cultures involving *Prochlorococcus* strains NATL2A, MED4 and MIT0604, culture decline was not monotonic, and was interrupted by additional growth phases about 40-50 and 100 days after the cultures started declining (Figure 4D). These latter growth phases were mostly absent in co-cultures with MIT9312 and MIT9313 (RMSE 0.2±0.1 vs 0.36±0.1 in the other strains, t-test p < 0.001). The correlation between N biomass and the secondary growth phases (i.e., higher deviation from simple Weibull model, Figure 4C) suggest that these phases may also be related to the ability of the interacting partners to recycle N through mutually-beneficial metabolic interactions.

### Conclusions and Future Prospects

Elucidating the mechanisms of microbial interactions requires well-characterized model systems. However, extending the insights from such models across the diversity of organisms and environmental conditions remains challenging. Our results from the highly-simplified system of multiple *Prochlorococcus* and *Alteromonas* strains provide an important step towards this goal. Using the rich information on interaction phenotypes present in the growth and decline curves, we identify conserved and strain-specific facets of these interactions. Despite the genetic diversity across the *Alteromonas* strains studied (30), it was primarily the identity of the *Prochlorococcus* strain that determined the interaction phenotype. This manifests in the growth and decline rates, in the shape of the curve (primarily the decline phase), in the amount of N retained in biomass and in whether the co-cultures are mutually synergistic or, potentially, competitive.

Under our laboratory conditions, it is likely that the combined response of both interacting partners to nitrogen starvation underlies the dynamics of the long-term co-cultures, although other stressors such as the increase in osmolarity/salinity or the accumulation of waste products cannot be ruled out (18,34,60). This response is dynamic, as illustrated by the reproducible deviations of the fluorescence curves from the monotonic decline predicted by all models tested (“second growth” stages, Figure 4). Three different (non-mutually-exclusive) processes may underlie these dynamics. Firstly, it is likely that one or both organisms modify their physiology or metabolism over time, for example through the activation of stringent responses, utilization of N or C storage pools, rewiring of metabolism to utilize available N sources or activation of mechanisms such as extracellular enzymes allowing the cells to access previously unusable substrates (e.g. (61,62)). Secondly, it is possible that there are “invisible” ecological dynamics underlying the observed fluorescence curves, for example cyclic changes in the abundance of *Alteromonas* cells. Under such a scenario, rapid *Prochlorococcus* mortality could produce an increase in *Alteromonas* abundance, resulting in degradation and remineralization of dead *Prochlorococcus* biomass and the release of resources that can drive subsequent *Prochlorococcus* growth. Finally, both *Prochlorococcus* and *Alteromonas* populations may be evolving, for example through emergence of genetically-distinct populations better adapted to nutrient starvation (reminiscent of the GASP phenotype described in *E. coli* and other bacteria (63)).

Why is it the identity of the primary producer (*Prochlorococcus*) rather than the heterotrophic “recycler” (*Alteromonas*) that determines the outcome of the co-culture? *A-priori*, it was reasonable to assume that the co-culture phenotype would be affected by the differences between the *Alteromonas* strains in their ability to degrade and utilize polysaccharides and a variety of other organic molecule (30,64). We speculate that the increased growth of *Alteromonas* in the co-cultures compared to the axenic ones is fueled primarily by the availability of major biomass components released by *Prochlorococcus* as they die, such as proteins, amino acids and nucleotides. Such common macromolecules do not require highly specialized metabolic processes to degrade and utilize, and hence can be utilized by all of the *Alteromonas* strains (65). It is possible that the differences between *Alteromonas* strains may manifest when more complex macromolecules are available, e.g. from plant material, or when all of the “easy to digest” (labile) organic matter has been utilized and only complex macromolecules remain (66). These conditions may not have been met in our experiments. Similarly, we currently do not know why some *Prochlorococcus* strains support a mutually-synergistic interaction with *Alteromonas* relatively early during the long-term N starvation (day 60) whereas other strains do not, and why at a later stage (day 100) almost all interactions are mutually beneficial. We could not identify any metabolic traits (11) clearly differentiating MIT9313 and MIT9312 (the “competitive” strains) from the others, suggesting more subtle differences exist between the *Prochlorococcus* strains in the organic matter they produce or in their response to N starvation (e.g. (67,68)).

Our results identify patterns in the interactions between clades of abundant marine phototrophs and heterotrophs, under conditions where nutrients are scarce, and their availability likely depends on recycling between phototrophs and heterotrophs. Whether or not such mechanisms may be physiologically relevant in the oligotrophic ocean, much of which is N-stressed (40), remains to be tested (conditions in laboratory batch cultures are very different from those in the nutrient-poor ocean). It is, however, noteworthy that the high heterotroph/phototroph biomass ratio observed during long-term N starvation here and in other studies (18) is similar to that of much of the open oligotrophic ocean (e.g. (69)) and references therein). Furthermore, *Alteromonas* may allow *Prochlorococcus* to adapt to light starvation (42) and to the presence of ROS (e.g. (70)), other stressors that can be encountered in the open ocean. However, the supportive role of *Alteromonas* cannot be taken for granted, as it also depends on culture conditions, for example CO_2_ concentrations (27).

Finally, the co-cultures did not reach a steady state, and did not represent a closed system. Thus, processes not represented in these simplified laboratory co-cultures, are necessary to explain the long-term stability over decades of *Prochlorococcus* in the oceans (71). Such processes could include multi-organism interactions, as natural communities are much more complex than the laboratory co-cultures, as well as oceanographic processes such as nutrient injection through deep mixing. More generally, cell mortality is intimately linked with the amount and type of recycled organic matter, yet the rate of mortality in natural communities is highly unconstrained (72). Hence, better representation of mortality in mathematical models (e.g. the use of appropriate mortality formulations) is likely important for understanding biogeochemical cycles (72). This study is a reminder that bacterial growth curves represent an information-rich representation of the life and death of cells, which can be mined to identify potential physiological changes of ecological relevance.

## Materials and Methods

### Strains and experiment set up

Axenic *Prochlorococcus* strains MED4 (HLI), MIT9312 (HLII), MIT0604 (HLII), NATL2A (LLI) and MIT9313 (LLIV) were maintained under constant cold while light (27 μmole photons m^−2^ s^−1^) at 22ºC (12,73). We used Pro99 media that was modified by reducing the concentration of NH_4_ from 800μm to 100μm (Pro99-LowN), resulting in *Prochlorococcus* entering stationary stage due to the depletion of available inorganic N (74). *Alteromonas* strains HOT1A3, Black sea 11, ATCC27126, AltDE1 and AltDE were maintained in ProMM media (43). Prior to the experiment, the axenicity of the *Prochlorococcus* cultures was tested by inoculating 1 ml culture into 15 ml ProMM (73). At the start of each co-culture experiment, *Alteromonas* cells from stationary-stage cultures (24-72 hour old) were centrifuged (10 minutes, room temperature, 10,000g), the growth media decanted, and the cells re-suspended in Pro99-LowN. The *Prochlorococcus* cultures (growing exponentially) and the re-suspended *Alteromonas* cells were then counted using BD FACSCanto™ II Flow Cytometry Analyzer Systems (BD Biosciences). For each co-culture, 1×10^6^ *Prochlorococcus* cells mixed with 1×10^7^ *Alteromonas* cells. For mono-cultures, 1×10^6^ and 1×10^7^ were added for *Prochlorococcus* and *Alteromonas*, respectively. The experiment was performed using triplicate 20ml cultures in borosilicate test tubes (2.5cm diameter, 15cm length).

### Fluorescence and Flow cytometry

Bulk chlorophyll fluorescence (FL) (ex440; em680) was measured almost daily using a Fluorescence Spectrophotometer (Cary Eclipse, Varian). Samples for flow cytometry were taken after 60, 100 and 140 days of experiment E1, fixed with glutaraldehyde (0.125% final concentration), incubated in the dark for 10 min and stored in -80 °C until analysis. 2 μm diameter fluorescent beads (Polysciences, Warminster, PA, USA) were added as an internal standard. Data was acquired and processed with FlowJo software. Flow cytometry was performed unstained to count *Prochlorococcus* cells followed by staining with SYBR Green I (Molecular Probes/ ThermoFisher) to count *Alteromonas* cells.

### Growth rate

Growth was computed - by solving the equation:

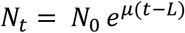

Where *N*_*t*_ represents the number of cells at time *t, N*_0_ is the initial number of cells, *µ* is the growth rate, and *L* is the growth lag. The LAN transformed equation was used to compute growth rate:

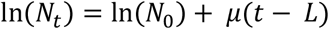

Linear regression was run on the growth phase, predicting ln(*N*_*t*_) based on time *t* with R^2^ > 0.9. The growth rate µ is the regression coefficient.

### Fit to Decline models

The following functions were used to fit against the measured fluorescence:

Exponential: 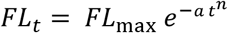

Bi-exponential: 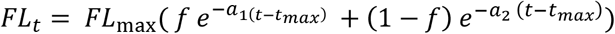

Harmonic: 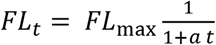

Weibull: 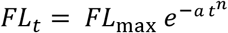

Where *FL*_*t*_ is the Fluorescence measured at time *t, FL*_*max*_ is the maximum fluorescence measured, *t*_*max*_ is the time when the fluorescence was highest, and *a, a1, a2, n, f* are the model parameters estimated by the fitting function.

The decline function was fit against each growth curve via curve_fit() function from scipy package (1.3.0), using the parameters: method=‘dogbox’, loss=‘soft_l1’, f_scale=0.1. Each model was fit using random initial parameter values and the initial values of 0.5 per parameter, and the fit with the lowest RSME selected. Goodness of fit was measures using root mean square error (RMSE). In the Weibull model the time needed to reduce the population by *d* factors of 10 (*t*_*d*2_) was estimated as in (59), using the formula:

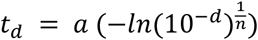

### Random Forest Classification

To detect difference in the curve pattern and not timing specific differences, the curves were aligned such that max growth point are at time zero, and time points from 10 days prior to max growth to 80 days after were selected. Since the specific measurement time points were different in different experiments and samples, rolling average was used to get mean fluorescence per day, and interpolation used to fill in missing measurements. The Fluorescence measurements were standardized via standard scalar by subtracting the mean and scaling to unit variance of each feature. Random Forest model was run in 10x cross validation to classify the curves by *Prochlorococcus* and by *Alteromonas* strains. To find the most significant days in *Prochlorococcus* classification, the model was built 30 times and the mean importance of all features (i.e., measurement days) calculated. Data preprocessing was done by pandas (0.25.0). Scaling and model fitting using sklearn (0.21.2).

### PCA ordination

PCA ordination was run on the growth curves. The fluorescence measurements were standardized via standard scalar by subtracting the mean and scaling to unit variance of each feature. Ordination was computed via principal component analysis (PCA). Data preprocessing was done by pandas (0.25.0). Scaling and PCA was done using sklearn (0.21.2).

### Statistics

Statistics were computed using the statsmodels package in python. Multi test correction was done by t_test_pairwise() using Bonferroni correction. Permanova analysis by adonis2 from R vegan package (R 3.61, vegan 2.5-7).

## Acknowledgements

We thank Sher lab members and especially Natalie Andrawes for help in fluorescence measurements and Daniel Segrè, Hans-Peter Grossart, Zhen Wu and Tal Luzzatto Knaan for critical reading of the manuscript. This work was supported by the Human Frontiers Science Program (grant RGP0020/2016, to DS) and the U.S.-Israel Binational Science Foundation (BSF) - U.S. National Science Foundation (NSF) program in Oceanography (grant 1635070/2016532, to DS). The funders had no role in study design, data collection and analysis, decision to publish, or preparation of the manuscript.

## Competing Interests

The authors declare no competing financial interests

## Notes

https://www.ncbi.nlm.nih.gov/sra/?term=PRJNA771345

### Competing Interest Statement

The authors have declared no competing interest.

