## supplemental text and figures for "Phototroph-heterotroph interactions during growth and long-term starvation across *Prochlorococcus* and *Alteromonas* diversity"

### Supplementary text S1 The differences between *Prochlorococcus* strains are most evident in the decline and long-term starvation stages

We used Random Forest Classification to test whether the observation that the co-culture outcome is determined by the *Prochlorococcus* and not *Alteromonas* strains, suggested by the clustering of the growth curves observed in the PCA ordination, is reproduced using an independent method. We then quantitatively determined which stages of the co-culture were most different between the *Prochlorococcus-Alteromonas* pairs. Random forest is a Supervised Machine Learning Algorithm widely used in classification and regression problems because it produces relatively accurate results while avoiding overfitting. Random forest has the added benefit that it provides feature importance, a measure of how much the model relies on each feature for classification. The algorithm accurately predicted, from the growth curves, the *Prochlorococcus* strain (10x cross validation accuracy 0.92±0.17), but performed much more poorly when trying to predict the *Alteromonas* strain (accuracy 0.62 ± 0.58) (Supplementary Figure S2A). This provides further support for the observation that the shape of the growth and decline curves was driven by the specific *Prochlorococcus* strain and not by the *Alteromonas*. The classification of the *Prochlorococcus* strains relied heavily on days during the decline phase where the dynamics of decline differed between strains. For example, around day 40 the decline rate of MIT9313 (LLIV) co-cultures seems to increase, whereas those of MIT9312 (HLII) were stable and the fluorescence of NATL2A (LLI) co-cultures actually increased (Supplementary Figure S2B).

### Supplementary text S2: The phenotype of interactions is maintained over multiple cycles of growth-decline-starvation

In all the subsequent transfers, the co-cultures declined relatively slowly, and were able to survive transfer for up to 140 days (Figure 1). The different *Prochlorococcus* strains could still be differentiated in a PCA ordination based on their growth curves (Adonis, p=0.001, Supplementary Figure S3), although the difference between low-light adapted NATL2A strain and the high-light adapted strains was less pronounced. Similar to E1, the clustering by *Alteromonas* strain was not as significant and r values were lower (Adonis, p > 0.1). These results are repeated in Random Forest classification. The accuracy of *Prochlorococcus* strain classification on subsequent transfers is 0.8-0.9, while the accuracy of *Alteromonas* classification is only 0.2-0.65 (Supplementary Figure S2A). Thus, the differences between *Prochlorococcus* strains in the way in which they interact with multiple *Alteromonas* strains (and the lack of any observed effect of the Alteromonas strains) are robust to the initial cell numbers and the time in co-culture.

Nevertheless, there were some consistent changes between E1 and all subsequent experiments. Firstly, the maximum fluorescence decreased in subsequent co-cultures compared to E1 (ANOVA, p<0.05, Supplementary Figure S4A). Secondly, the growth rate of *Prochlorococcus* in co-culture increased in most strains and experiments (ANOVA, p<0.05, Supplementary Figure S4C). Thirdly, MIT9313 (LLIV), which was inhibited in the first co-culture by all Alteromonas strains, did not show this phenotype in subsequent transfers (Figure 1A, Supplementary Figure S4B). This is potentially due to the lower number of inoculated Alteromonas cells, as shown previously (1).

### Supplementary text S3: Carrying Capacity of the cultures

The carrying capacity of the cultures was defined as the amount of nitrogen retained in cell biomass (rather than as dissolved organic N) at various stages of long-term co-culture. Cell numbers from flow cytometry were converted into nitrogen using 7 fg N cell$^{-1}$ for the high-light strains MED4, MIT9312 and MIT0605, 10.5 fg N cell$^{-1}$ for strain NATL2A and 14 fg N cell$^{-1}$ for strain MIT9313 (2,3). For Alteromonas we used a value of 13 fg N cell$^{-1}$ (4,5). We note that these are the values we used are at the lower end of measured cell values, which reach up to 20 fg N cell$^{-1}$ for low-light *Prochlorococcus* and 25 fg N cell$^{-1}$ for *Alteromonas*, since using the higher N cell quota leads to biomass that is higher than the total nitrogen available in the system (see below). This assumption is supported by studies showing that cells contain less nitrogen under long term N stress compared to exponential growth (6–8).

The cell numbers were converted to µmol/L by the formula:

biomass [µmol/L] = N [cell/ml] * $Q_N^{PRO}$ [fg/cell] * 1e-9 [converting femtomol-> micromol] / 1e-3 [ml->L] / $MW_N$ [g/mol]

Where N is the number of cells per ml, $Q_N$ is the cell N quota, and $MW_N$ is the molecular weight of nitrogen.

Since neither *Prochlorococcus* nor *Alteromonas* are known to fix nitrogen or perform denitrification, we assume that the co-cultures are closed systems for N (i.e. the total N in the system does not change over time). The experiment media contains 100 µmol / L of $NH_4$ (DIN), an estimated 5 µmol/L of DON (9) from the natural seawater used for the media, and the N biomass of the inoculated cells. The initial N in the media is calculated as:

initial_N = 100 [DIN] + 5 [DON] +

$Q_N^{PRO}$ [fg/cell] * 1e6 [cell/ml] * 1e-6 [f->u, ml->L] / 14 [$MW_N$] +

$Q_N^{ALT}$ [fg/cell] * 1e7 [cell/ml] * 1e-6 [f->u, ml->L] / 14 [$MW_N$]

### Supplementary Tables

#### Table S1: N carrying capacity

| | Available N [µmol N/L] | Total N Biomass [µmol N/L] | | |
|---|---|---|---|---|
| | | **Co-culture** | ***Alteromonas* Only** | ***Prochlorococcus* Only** |
| **N** | | 71 | 13 | 14 |
| **Day 0** | DIN: 100<br>DON: 5<br>PRO: 0.7<br>ALT.: 18<br>Total: 123.6 | | | |
| **Day 60** | | PRO: 16±11<br>ALT: 54±29<br>Total: 69±35<br>% of available N: 56±28% | 32±22<br>% of available N: 26±18% | 0.01±0.00<br>% of available N: 0% |
| **Day 100** | | PRO: 6±4<br>ALT: 18±8<br>Total: 23±10<br>% of available N: 19±8% | 4±4<br>% of available N: 3±3% | 0.03±0.12<br>% of available N: 0% |
| **Day 140** | | PRO: 0.5±0.8<br>ALT: 0.8±0.7<br>Total: 1.3±1.1<br>% of available N: 1±1% | Not measured | 0<br>% of available N: 0% |

* PRO: *Prochlorococcus*, ALT: *Alteromonas*

**Table S2: Decline model data**

| | | E1 Axenic (n=13) | E1 Co-culture (n=73) | E2 Co-culture (n=72) | E3 Co-culture (n=65) | E2.1 Co-culture (n=69) | E2.2 Co-culture (n=64) |
|---|---|---|---|---|---|---|---|
| **Weibull shape** | | 2.10±0.88 | 0.43±0.17 | 0.48±0.32 | 0.48±0.40 | 0.65±0.34 | 0.53±0.16 |
| **RMSE** | Exponential | 0.26±0.11 | 0.51±0.30 | 0.21±0.10 | 0.22±0.12 | 0.24±0.14 | 0.23±0.14 |
| | Biexponential | 0.26±0.11 | 0.27±0.17 | 0.13±0.07 | 0.12±0.09 | 0.13±0.06 | 0.14±0.09 |
| | Harmonic | 0.50±0.19 | 0.32±0.20 | 0.15±0.08 | 0.14±0.07 | 0.16±0.09 | 0.17±0.09 |
| | Weibull | 0.14±0.08 | 0.29±0.13 | 0.16±0.09 | 0.13±0.07 | 0.15±0.08 | 0.16±0.08 |
| **BIC** | Exponential | -231±80 | -116±82 | -231±78 | -117±45 | -221±87 | -142±65 |
| | Biexponential | -222±80 | -208±79 | -296±92 | -157±50 | -296±77 | -176±65 |
| | Harmonic | -122±81 | -187±77 | -274±79 | -151±48 | -275±87 | -166±64 |
| | Weibull | -331±97 | -188±57 | -268±90 | -151±45 | -279±79 | -167±58 |

### Supplementary Figures

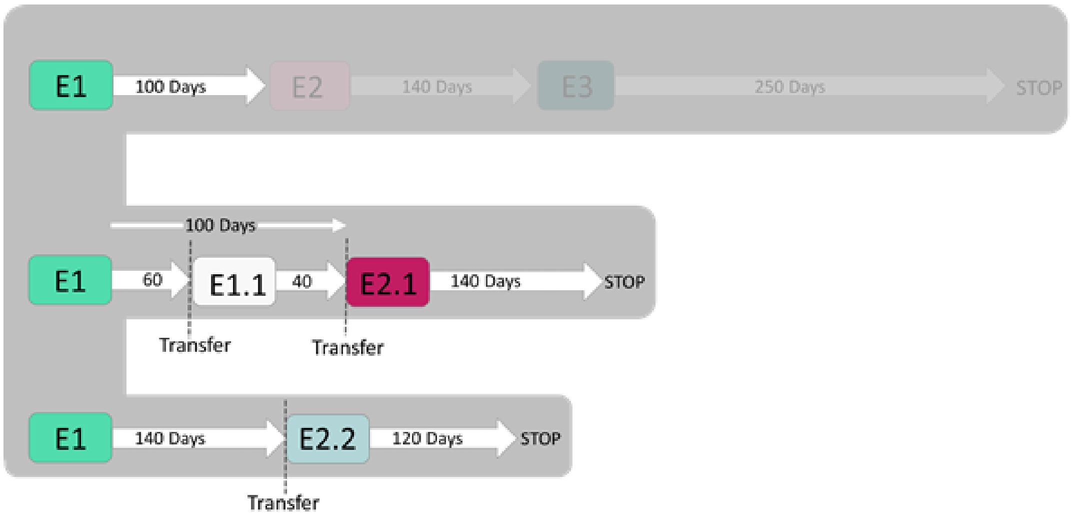


**Figure S1: Additional Independent transfers from E1.** Three separate transfers were performed from the first experiment (E1). The top line represents experiments 2 and 3, which are the focus of the study and are shown in Figure 1B, C. Two additional transfers were performed, further supporting some of the observations described in the main text. The middle line shows two short transfers from E1 (after 60 and then 40 days, E1.1), to assess the viability of the axenic cultures. These were then transferred subsequently to E2.1, which was maintained for 140 days. The lower line shows an additional transfer from E1 after 140 days to E2.2, which was maintained for another 120 days. E2.1 therefore represents a culture transferred after relatively short N starvation, whereas E2.2 represents an independent replicate of transfer after long-term starvation (similar to E2 but after longer starvation).

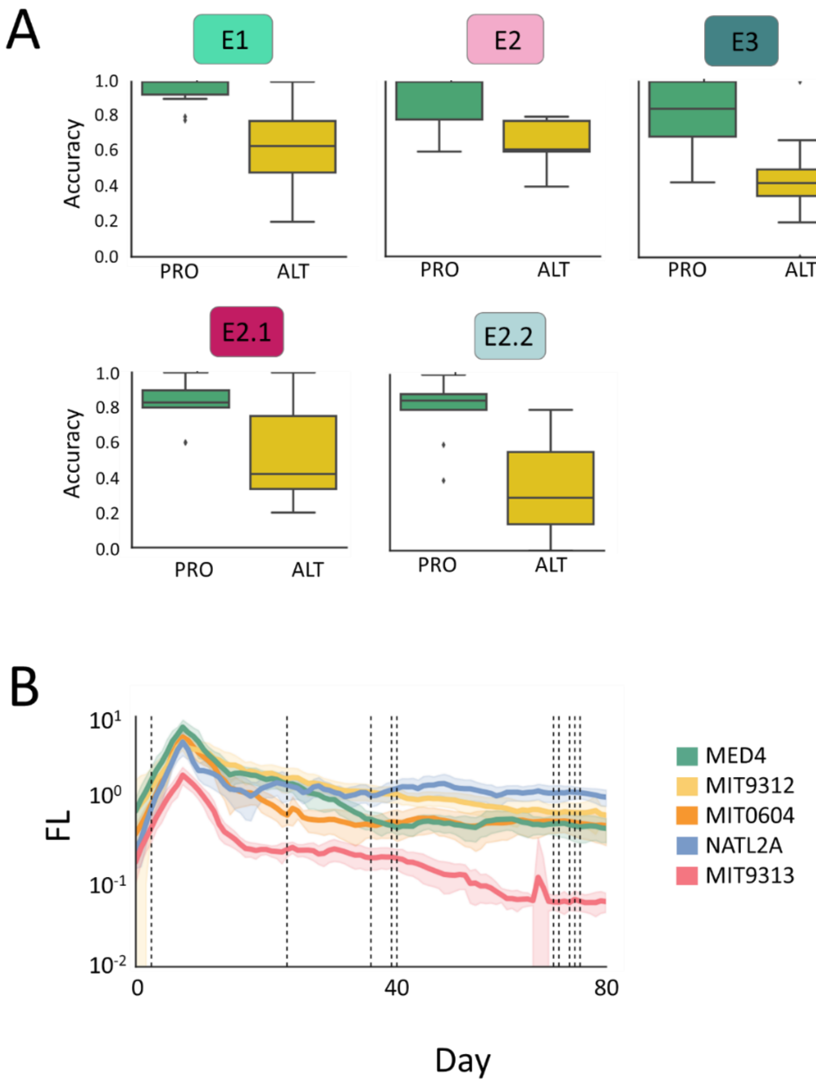


**Figure S2: Classification of *Prochlorococcus* and *Alteromonas* strains and identification of important culture stages differentiating between strains using machine learning.** A. Random Forest accuracy of 10x cross validation when predicting *Prochlorococcus* (PRO) and *Alteromonas* (ALT) classification based on their growth curves. The algorithm has higher cross-validation accuracy in predicting *Prochlorococcus* strain in all experiments/transfers, suggesting that the *Prochlorococcus* strain, and not the *Alteromonas* one, has a stronger effect on the shape of the growth and decline curves. The error bars represent 10 model builds and their respective cross-validation runs. B. Significant days in the Random Forest classification of the ***Prochlorococcus-Alteromonas*** co-culture curves in E1. The days with the 10 highest feature importance are marked. All curves were aligned to max growth day.

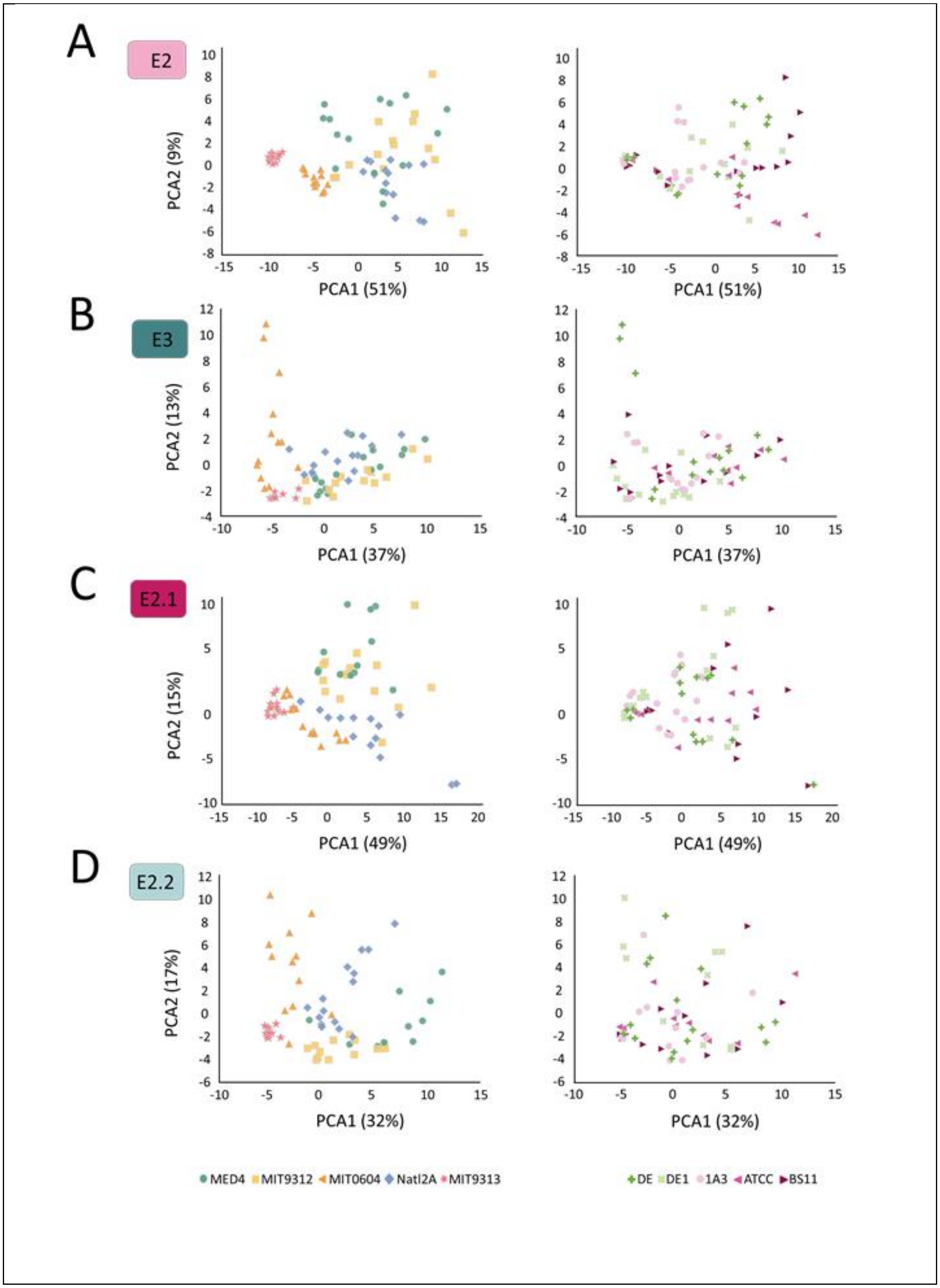
A
E2
B
E3
C
E2.1
D
E2.2
PCA1 (51%)
PCA2 (9%)
PCA1 (37%)
PCA2 (13%)
PCA1 (49%)
PCA2 (15%)
PCA1 (32%)
PCA2 (17%)
MED4
MIT9312
MIT0604
Natl2A
MIT9313
DE
DE1
1A3
ATCC
BS11

**Figure S3: PCA of the fluorescence curves from subsequent experiment.** In all experiments, the curve shapes clustered primarily based on the *Prochlorococcus* and only to a much lower extent by the *Alteromonas* strains in co-culture.

Adonis2:

A. E2: *Prochlorococcus* $F(4,67) = 13.02$, $p = 0.001$. *Alteromonas* $F(4,67) = 3.18$, $p = 0.001$

B. E3: *Prochlorococcus* $F(4,60) = 8.85$, $p = 0.001$. *Alteromonas* $F(4,60) = 1.96$, $p = 0.02$

C. E2.1: *Prochlorococcus* $F(4,68) = 14.07$, $p = 0.001$. *Alteromonas* $F(4,68) = 2.79$, $p = 0.001$

D. E2.2: *Prochlorococcus* $F(4,59) = 10.08$, $p = 0.001$. *Alteromonas* $F(4,59) = 0.72$, $p = 0.8$

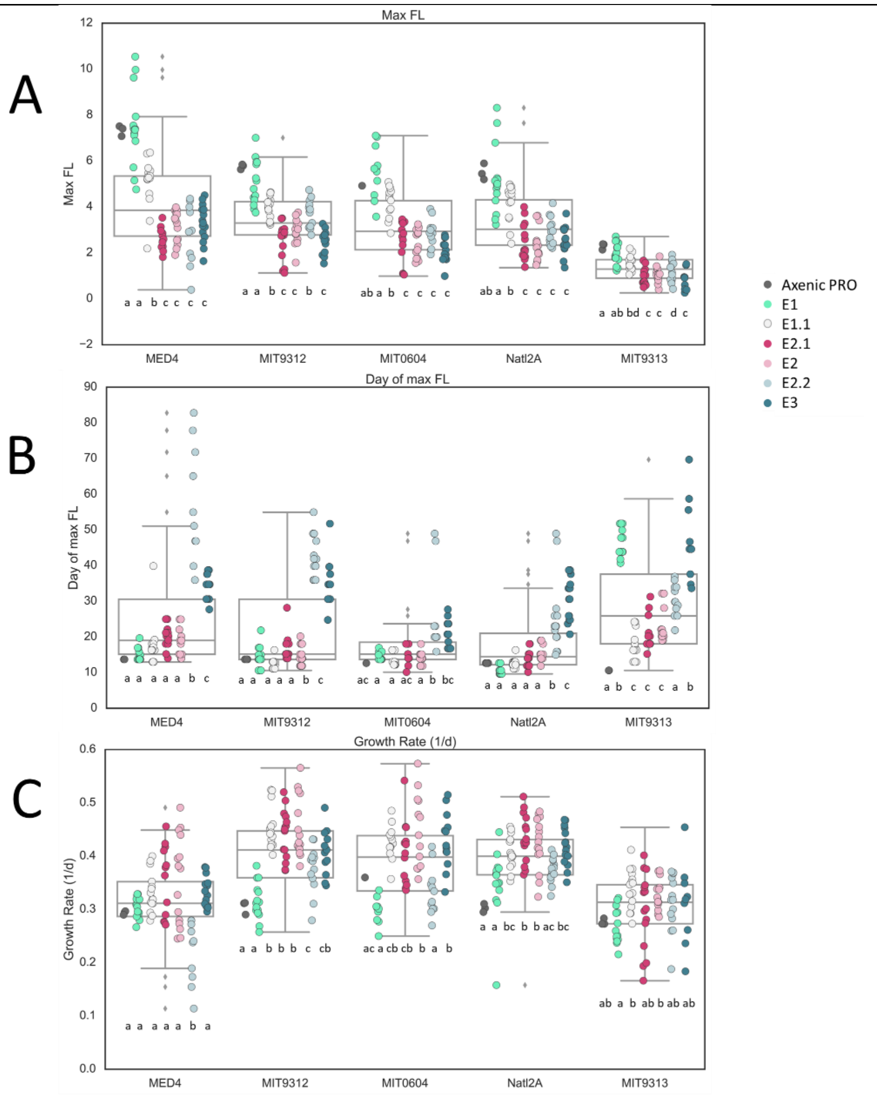


**Figure S4: Growth and growth rate of subsequent co-cultures.** In all panels, characters indicate results of ANOVA with Bonferroni correction between the different transfers per *Prochlorococcus* strain (different characters indicate $p < 0.05$). A. Maximum Fluorescence (proxy for number of cells) B. Day of max Fluorescence. c. Growth rates (Axenic PRO: Axenic *Prochlorococcus* in E1). In the subsequent co-cultures (after E1), the growth rate of *Prochlorococcus* increased compared to the first interaction between *Prochlorococcus* and *Alteromonas.* One exception was E2.2, in which growth rates were similar to E1 or (in the case of MED4) even lower. We currently have no explanation for this observation.

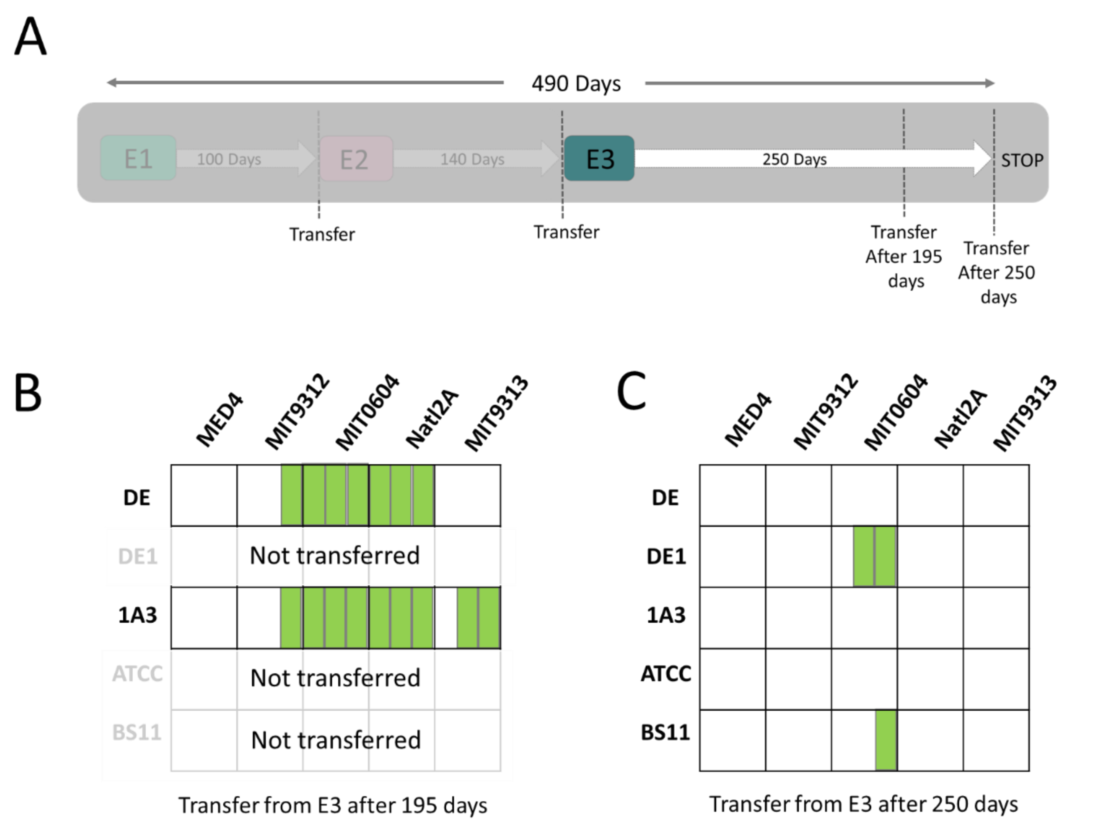


**Figure S5: Co-cultures able to survive for extended period**. A. Co-cultures from E3 were transferred to new media at two time-points (195 and 250 days). B. On day 195, only co-cultures with *Alteromonas macleodii* HOT1A3 and *A. mediterranea* DE were transferred (a total of 30 cultures), of which 16 survived transfer (green squares). C. After 250 days all 75 co-cultures were transferred into fresh media, with only three surviving transfer.

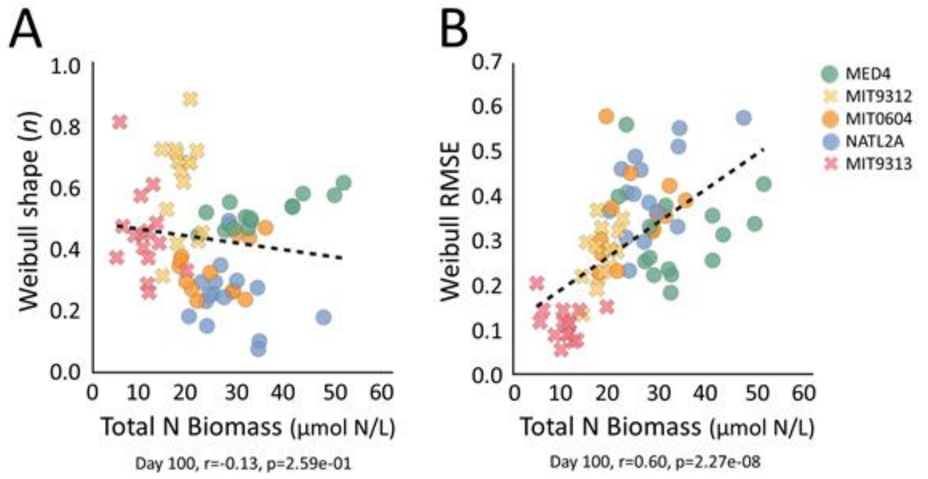


**Figure S6: Correlations between total N biomass on day 100, the Weibull shape (A) and the Weibull RMSE (B).** Pearson's r and p values are shown below the plots.
